# Neural correlates of auditory pattern learning in the auditory cortex

**DOI:** 10.1101/2020.09.24.311464

**Authors:** H. Kang, R. Auksztulewicz, H. J. An, N. Abichacra, M. L. Sutter, J. W. H. Schnupp

**Affiliations:** Department of Neuroscience, City University of Hong Kong; Neuroscience Department, Max Planck Institute for Empirical Aesthetics, Frankfurt, Germany; Center for Neuroscience, and Section of Neurobiology, Physiology and Behavior, University of California, Davis, California

**Keywords:** auditory perception, learning, electrocorticography, rat, auditory cortex

## Abstract

Learning of new auditory stimuli requires repetitive exposure to the stimulus. Fast and implicit learning of sounds presented at random times enables efficient auditory perception. However, it is unclear how such sensory encoding is processed on a neural level. We investigated neural responses that are developed from a passive, repetitive exposure to a specific sound in the auditory cortex of anesthetized rats, using electrocorticography. We presented a series of random sequences that are generated afresh each time, except for a specific reference sequence that remains constant and re-appears at random times across trials. We compared induced activity amplitudes between reference and fresh sequences. Neural responses from both primary and non-primary auditory cortical regions showed significantly decreased induced activity amplitudes for reference sequences compared to fresh sequences, especially in the beta band. This is the first study showing that neural correlates of auditory pattern learning can be evoked even in anesthetized, passive listening animal models.

## Introduction

Sensory perception requires correctly recognizing incoming sensory stimuli by extracting relevant information from memory. Such memory is formed by implicit learning of sensory input through repetitive exposure. Fast memory formation by capturing unique features of sensory signals is thus one key factor for efficient sensory perception.

In hearing, a series of recent studies reported fast and robust learning of abstract sounds, using a novel experimental paradigm that resembles unsupervised implicit learning of newly presented acoustic stimuli in natural scenes (Agus et al., 2010; Andrillon et al., 2015; Luo et al., 2013). In this paradigm, participants were simply asked to detect a within-sequence repetition in random noise samples. Unbeknownst to them, one specific noise sample would re-occur occasionally, and even though the subjects were unaware of this, they nevertheless showed fast, selective improvement in processing the frozen “reference” stimulus, which implies a rapid and robust memorization of random features of complex sounds. Such behavioral improvement for the re-occurring sound was supported by increased inter-trial coherence of brain responses for the re-occurring stimulus compared to other random stimuli measured by subsequent EEG and MEG studies in humans (Andrillon et al., 2015; Luo et al., 2013). Interestingly, increases in neural coherence could even be observed when the human subjects were in Rapid Eye Movement (REM) or light non-REM sleep during the experiment (Andrillon et al., 2017), suggesting that a neural index related to learning new sounds can be traced even following passive exposure. While these findings provided insights into the neural correlates of implicit learning of new auditory stimuli, further investigations using invasive measurements will be needed to understand the underlying mechanisms. The present study aimed at investigating neural responses shaped by passively presented re-occurring sounds in the auditory cortex using rats as an animal model.

Previous electrophysiological studies have investigated how neurons adapt to re-occurring sounds to understand memory and adaptation processes, by using a simplified experimental paradigm, in which a series of standard sounds (usually pure tones) is disrupted by presentation of a deviant sound (Garrido et al., 2009; Malmierca et al., 2014; Nieto-Diego and Malmierca, 2016). Under such paradigms, stimulus specific adaptation (SSA) has been widely reported using comparisons between habituated neural responses to a standard sound against the typically greater responses for a novel, deviant sound. SSA effects have been observed along the auditory pathway, first in the primary auditory cortex (AC), and then also in non-lemniscal subdivisions of the inferior colliculus (IC) and the medial geniculate body (MGB; Anderson et al., 2009; Ayala and Malmierca, 2013; Parras et al., 2017; Ulanovsky et al., 2003). A more recent study further reported stronger SSA in non-primary AC fields compared to primary AC (Nieto-Diego and Malmierca, 2016). Another study using more complex and realistic sounds has suggested that higher-order regions in the AC, rather than primary fields, may be uniquely susceptible to adaptation to repeatedly presented realistic auditory inputs (Lu et al., 2018). The study further reported that the adaptation effect was retained after the disruption period from another repetitive presentation of the other sound input in the AC. These results point to the active involvement of the AC in learning and adaptation to ongoing or predictable sounds, which is thought to play a role not only in encoding stimuli, but also their context (Bar-Yosef et al., 2002; Lu et al., 2018; Skipper, 2014). However, while previous studies compared neural responses evoked by occasional deviants relative to consecutively presented standards, such constant presentation of a single sound is rare in realistic auditory scenes. Instead, recognizable re-occurring sounds typically appear occasionally, interspersed with other random, non-repeating sounds, and yet listeners learn them without much effort.

In the present study, instead of the classical paradigm of constant representations of a single sound, we adapted an experimental paradigm (Agus et al., 2010) to intermittently present frozen “reference” sequences among other random sequences. The aim of the present experiment was to look for a physiological correlate of the “learning” of the frozen sequence that can occur even during passive exposure in the AC of anesthetized rats, using electrocorticography (ECoG) as a first step for identifying neurophysiological markers. We focused on investigating neural characteristics that emerged by learning re-occurring auditory patterns across primary and non-primary auditory fields within the AC. We particularly controlled for physical differences between stimuli by comparing pure induced neural responses, non-phase locked responses to the stimulus computed after the time-frequency decomposition rather than evoked neural responses. By doing so, we could minimize any observed effect to be drawn from characteristics of the stimulus itself and focus on the neural modulations induced by higher-order processing (David et al., 2006; Klimesch et al., 1998). Our results show more attenuated induced activity amplitudes for the re-occurring sounds compared to other sounds, in both primary and non-primary fields, especially in the beta frequency band.

## Methods

### Participants

Six female adult Wistar rats (age = 8 −21 weeks, mean = 12.5, std = 4.42, weight = 257 – 315 g) were acquired from the Chinese University of Hong Kong. Experimental procedures were approved by the City University Animal Research Ethics Sub-Committee and conducted under license by the Department of Health of Hong Kong [Ref. No. (19-31) in DH/SHS/8/2/5 Pt.5].

### Stimuli

We generated sequences of acoustic stimuli shown schematically in Fig. 1A. Each sequence consisted of five segments. Sequences could be made up either of 0.24 s long dynamic random chords (DRCs) or of 0.2 s long white noise (WN) snippets. Sequences either consisted of the same segment repeated five times (repeated sequence, RS), or they were non-repeating, random sequences (S). To make it easier to distinguish neural signatures of repetition detection from simple onset or offset responses, sequences were prefixed and suffixed with additional “head” and “tail” segments, which were always generated afresh, and ramped on or off linearly. Segments were joined with 5 ms ramping overlaps to avoid transients. Sequences were presented in blocks at a rate of one sequence every 0.6 s. One block contained 100 unique RS and S sequences each, as well as one “frozen RS” and one “frozen S” sequence, which were presented 100 times each in each block. Adopting the nomenclature of Agus et al. (2010) we refer to the frozen sequences as “references”, or “RefRS” and “RefS” respectively. Thus one block consisted of a shuffled series of 400 stimuli, with 100 different S and RS sequences and 1 unique RefS and RefRS sequence being presented in random order (see Fig. 1B). For each block, sequences were generated anew with new random seeds.

**Figure 1.**
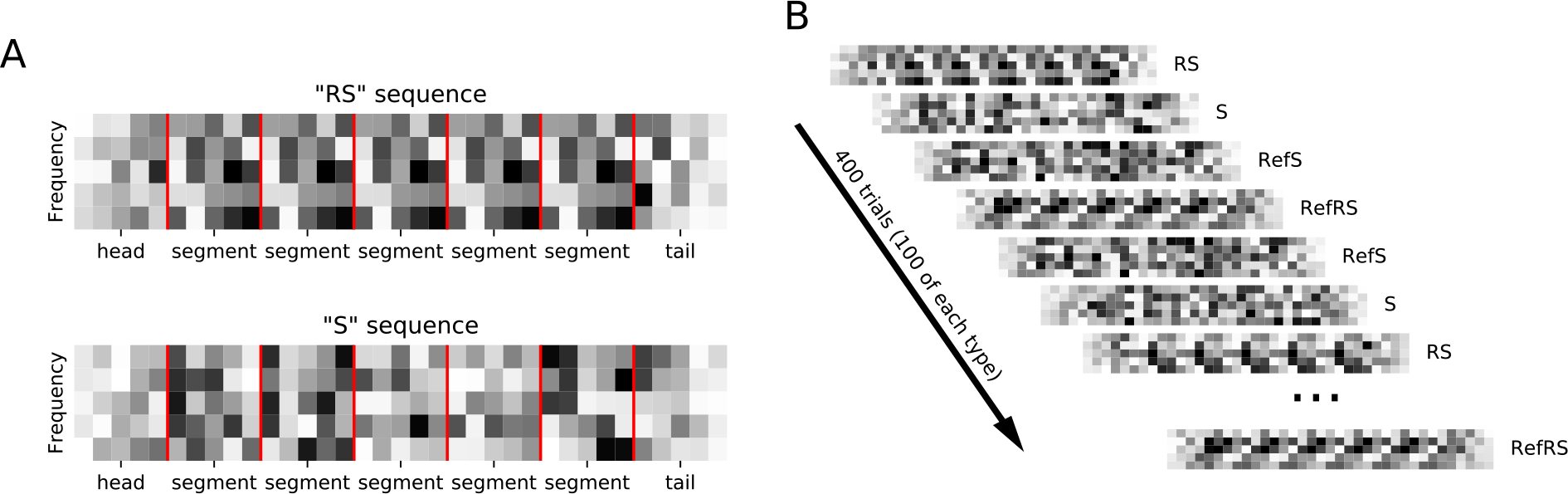
A: Sequences were composed of random spectral pattern (DRC or white noise) segments (marked by red vertical lines) which were either repeated 5 times in a row (RS sequence) or non-repeating (S sequence). Ramped, random “head” and “tail” segments bracketed each sequence. B: sequences were presented in blocks of 400 trials. Each block contained 100 sequences, each of unique R and S sequences as well as repeated “reference” RefRS and RefS sequences, which were presented in random order.

DRC sequences consisted of 12 chords of superimposed 20 ms pure tones at 15 log-spaced frequencies from 500 to 20,000 Hz. The level of each tone was randomly drawn from a uniform 50-90 dB SPL range to have mean 70 dB SPL, generating random spectro-temporal patterns characteristic of each DRC. The WN sequences consisted of Gaussian noise snippets, generated from different random seed values. As DRC segments comprise more salient spectral contrasts than WN segments, we expected within-sequence repetition to be easier to detect in DRC than in WN sequences.

### Experimental procedure

We recorded responses to five blocks of DRC sequences and five blocks of WN sequences from ECoG arrays placed onto the right auditory cortex (AC). Anesthesia was induced using Ketamine (80 mg/kg) and Xylazine (12 mg/kg, Intraperitoneal injection; i.p.) and maintained with Urethane (20%, 7.5 µl/g, i.p.). Urethane anesthesia minimizes NMDA receptor blockage and closely resembles REM and stage II nREM sleep-like status (Pagliardini et al., 2012). Dexamethasone (0.2 mg/kg, i.p.) was injected to prevent inflammation. Adequate anesthesia was confirmed by regular testing for the suppression of the toe pinch withdrawal reflex. Body temperature was kept at 36 ± 1 °C with a heating pad. The rat was placed in a stereotaxic frame and the head was fixed with hollow ear bars to allow the delivery of auditory stimuli. We measured the auditory brainstem responses (ABRs) in each ear to confirm that the rats had normal hearing sensitivity (click thresholds < 20 dB SPL). The right AC was exposed by a rectangular 5 × 4 mm craniotomy which extended from 2.5 to 7.5 mm posterior from Bregma, with its medial edge 4 mm from the midline. A 61-channel ECoG array (Woods et al., 2018) was connected to a Tucker Davis Technologis (TDT) PZ5 neurodigitizer and RZ2 real-time processor, and placed on the exposed cortex. The sound sequences were presented via a TDT RZ6 multiprocessor at a sampling rate of 48,828 Hz, and ECoG responses were recorded at 24,414 Hz using BrainWare software.

### Data analyses

Acquired neural responses were pre-processed to obtain event-related potentials (ERPs) for each channel and condition for each rat. ERPs were used to look for differences across the four conditions (S, RS, RefS, RefRS), using the time-frequency analysis described below. To calculate ERPs of each channel, ECoG signals were low-pass filtered (2^nd^ order zero-phase Butterworth) at 45 Hz, downsampled to 1,000 Hz, and re-referenced to the common mean. Time points at which signal values exceeded ± 3 std of the mean signal across time were identified as outliers and removed (i.e. replaced by linear interpolation from neighboring points, and detrending as described in (de Cheveigné and Arzounian, 2018). Signals were then epoched from −100 ms to 1600 ms relative to the onset of each sequence. Epochs for each condition were averaged to compute mean ERPs for each channel. To reduce data dimensionality, as well as minimize the effect of individual variations in electrode placement between rats, we subjected each rat’s channel-by-time ERP matrix (averaged across conditions) to a principal component analysis (PCA) and ordered components from the highest to the lowest amount of variance. We selected the top components (in order of variance explained) describing at least 99% variance and calculated the weighted sum of the spatial components to quantify the evoked response topography with reduced variabilities across rats. A visual inspection of regional response differences per rat from the obtained topography revealed that channels with the lowest response weights were mainly around A1 areas while channels with the highest response weights were mainly around non-A1 areas. Thus, we grouped top response-weighted channels as a tentative non-A1 cluster, and the bottom response-weighted channels as a tentative A1 cluster for further analysis of regional differences. Since the number of channels included in each group did not affect the result, we grouped the channels into the top 30 channels for the non-A1 cluster and the rest for the A1 cluster.

Next, to characterize the differences in induced responses to reference sequences (RefRS and RefS) compared to fresh sequences (RS and S), we ran a time-frequency analysis of single-trial ECoG signals using Morlet wavelets implemented in the FieldTrip toolbox for Matlab (frequency range: 4 - 80 Hz in 2 Hz steps; 400 ms fixed time window; Billig et al., 2019). The time-frequency power spectrum of each trial was rescaled by subtracting the time-frequency spectrum of the average ERP for the same condition (i.e., evoked power) on a logarithmic scale. This subtraction yielded an estimate of induced activity amplitude, whereby the responses in each individual trial did not have to be precisely time-locked to the stimulus (Hartmann et al., 2012). Therefore, the induced response differs from the ERP by focusing on the oscillation of spectral power rather than on phase-locked responses to the stimuli. The resulting single-trial induced responses were log-scaled and averaged across trials. After obtaining average time-frequency power spectra for each rat, channel group, and stimulus condition, we ran two cluster-based permutation paired *t*-tests (as implemented in FieldTrip) on RefRS vs. RS stimuli and on RefS vs. S, with 1000 iterations per test.

## Results

First, a cluster-based permutation paired *t*-test with 1,000 iterations revealed no significant differences of ERP amplitudes (averaged across channels) between RefRS and RS conditions or RefS and S conditions, either for DRC or for WN sequences. For both DRC and WN stimuli, channels presumed to be primary auditory cortex (A1) showed lower evoked response weights (averaged across all trials and conditions) than channels presumed to be non-primary auditory cortex, mainly from around suprarhinal auditory field (SRAF; Fig. 2A). Based on the evoked response weights, we grouped channels into two clusters, A1 and non-A1 clusters, based on the evoked response weights for further analyses on comparing induced activity amplitudes differences across conditions. A time-frequency analysis of induced activity revealed robust differences between pairs of Ref and non-Ref conditions for both clusters for DRC, but not for WN. We first focused on neural activity in the non-A1 cluster induced by DRC stimuli, based on the hypothesis that perceptual learning of complex stimuli may primarily modulate activity in higher-order regions. When comparing time-frequency responses between RefRS and RS conditions, we observed significantly decreased power for RefRS vs. RS during the sequence presentation, emerging from the onset of RefRS mostly in the beta band (10 – 40 Hz; T_min_ = −19.41, T_max_ = −2.57, all cluster-based *p*’s < 0.05). This was especially pronounced during the first three segments of the sequences (Fig. 2B). In the RefS vs. S comparison, decreased power for RefS was also observed from the RefS onset to the sound offset across the theta, alpha, and beta band (4 – 30 Hz; T_min_ = −8.54, T_max_ = −2.58, all cluster-based *p*’s < 0.05). In the A1 cluster, power decrease for RefRS vs. RS was observed in a similar frequency range (4 – 30 Hz; T_min_ = −19.82, T_max_ = −2.58, all cluster based *p*’s < 0.05) to the non-A1 cluster, but persisted for a longer time period (from sequence onset to sound offset), mostly in the beta band. Power decrease for RefS vs. S was observed for similar time period and frequency bands to the non-A1 cluster (4 – 30 Hz; T_min_ = −9.98, T_max_ = −2.57, all cluster-based *p*’s < 0.05). For WN sequences, we did not observe any significant differences in time-frequency response spectra between either RefRS vs. RS or RefS vs. S comparisons.

**Figure 2.**
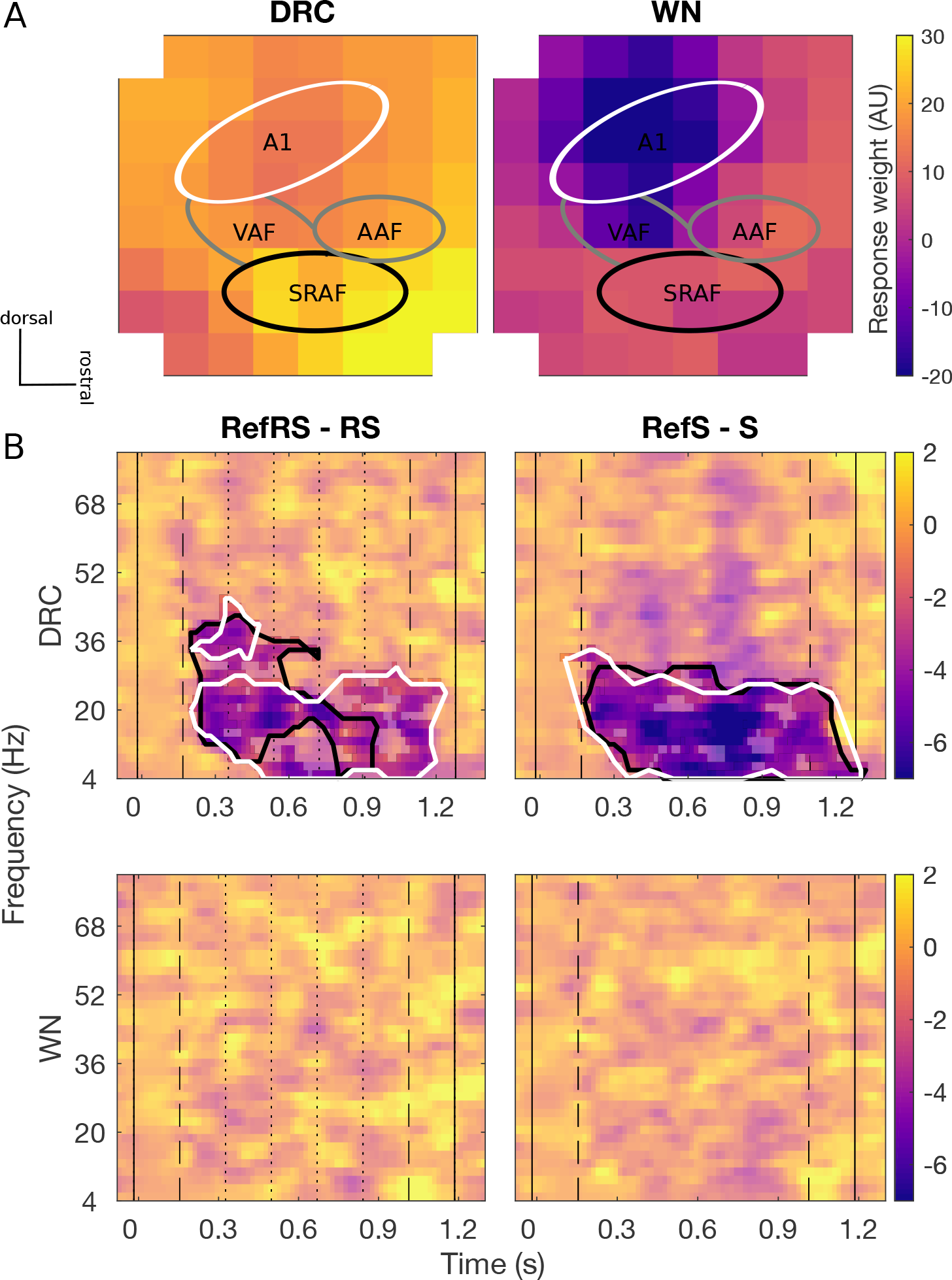
A: Spatial topography maps (methods of Nieto-Diego & Malmierca, 2016) of average evoked responses collected by 8 x 8 ECoG for DRC (left) and WN (right) sequences respectively. Colorscale is fixed for both sound types. Tentative subfields of the AC are marked with relevant labels (A1, VAF, AAF, and SRAF). The main A1 cluster (white line) shows slightly lower evoked response weights, and the putative SRAF cluster (black line) showed generally greater evoked response weights. Overall evoked responses to DRC sequences were stronger than for WN. In both cases, greater evoked responses were generally found in the non-A1 clusters. B: Differences in average time-frequency induced power spectra for RefRS minus RS (left) and RefS minus S (right) pairs for DRC (top) and WN (bottom). Black solid vertical lines indicate sound onset and offset, dashed vertical lines indicate reference sequence onset and offset, and dotted vertical lines on the left panel indicate within-sequence segment boundaries. Black contours in DRC spectra indicate time-frequency areas where a significant difference between conditions was observed for the non-A1 cluster, and white contours indicate the areas with a significant difference observed for the A1 cluster (cluster-based *p* < 0.05). No significant power difference was observed for both pairs in WN.

## Discussion

We assessed distinct neural correlates of implicit learning processes through a repetitive passive exposure to a specific auditory sequence. We compared neural dynamics of re-occurring sequences with the same acoustic characteristics (RefRS and RefS) and a group of other sequences that were presented only once (RS and S), by computing induced activity amplitude of neural signals recorded from primary (A1) and higher-order auditory cortex. We observed decreased induced activity amplitude throughout the stimulus sequence for RefRS and RefS compared to RS and S, mainly in the beta band, both for A1 and non-A1 channel clusters, but only for DRC stimulus sequences which contain more salient acoustical features compared to WN. This finding suggests an active involvement of both primary and non-primary AC in the implicit learning of complex auditory patterns.

Unlike most previous studies that computed differences between evoked responses as an index of learning (Lim et al., 2016; Lu et al., 2018), we did not observe any significant difference in ERPs across conditions. This result, however, was expected in our study as Ref sequences were presented in a passive listening setting with a complex and unpredictable experimental design. Previous neuroimaging studies in humans under similar paradigms also mainly focused on comparing inter-trial coherence rather than ERP differences between RefRS and RS or RefS and S (Andrillon et al., 2015; 2017; Luo et al., 2013). Thus, we focused on comparing induced response power obtained from each trial after the time-frequency decomposition. We found distinct response patterns for both of re-occurring sequences (RefRS and RefS) from the beginning of the sequence presentation when compared to fresh sequences (RS and S) for DRC. Such difference suggests that within-sequence repetitions are not a requirement for the learning process, as long as the sequence contains salient information to be learned. Both effects in our study were observed mostly in the beta band, which has been implicated in sensory memory (Haenschel et al., 2000; Scholz et al., 2017) and sensory predictions (Auksztulewicz and Friston, 2016; Auksztulewicz et al., 2017; Pearce et al., 2010).

The effect of attenuated induced activity amplitude for RefRS over RS and RefS over S was found from both A1 and non-A1 channel clusters. The effect mostly overlapped between the two clusters, especially for RefS. Interestingly, for RefRS, significant power differences in the non-A1 cluster started to diminish already after the first three segment repetitions – unlike the power difference in the A1 cluster which lasted towards the end of the sequence (Fig. 2B). Although further investigation is required, we hypothesize that repeated segments within RS may also have been learned, resulting in no distinctive difference between RefRS and RS to be observed towards the end of the sequence. This could further indicate that acoustic features presented in re-occurring brief segments that are as short as 200 ms can be effectively learned and recognized. Such characteristic was only observed in the non-A1 cluster, suggesting a hierarchical structure of the auditory cortex and indicating a role of higher-order regions in repetition suppression and prediction in a shorter time frame (Auksztulewicz and Friston, 2016; Lim et al., 2016; Lu et al., 2018). The present finding provides further insights into neural responses mediating RefRS learning within a short timeframe.

Finally, although a previous study using the similar paradigm showed neural correlates of RefRS using WN as stimuli (Andrillon et al., 2015; Luo et al., 2013), we did not observe any distinctive characteristic of RefRS and RefS for WN. This could be due to a greater saliency that DRC could generate for its recognition compared to WN, when such short repeating segment was presented. Other factors such as the segment duration or seamless presentation could also have affected the saliency of the Ref sequence (Agus et al., 2010; Andrillon et al., 2017), which requires further investigation.

The present experiment was conducted under anesthesia. Anesthetics could affect certain aspects of auditory processing such as spike timing, population activity or frequency tuning, depending on the type of anesthetics (Gaese and Ostwald, 2001; Huetz et al., 2009; Noda and Takahashi, 2015; Zurita et al., 1994). However, a large amount of previous physiology research investigating neural adaptation for re-occurring sounds or information processing has been conducted on animals under anesthesia (e.g., Anderson et al., 2009; Astikainen et al., 2011; Bao et al., 2004). These studies, as well as our results, suggest that neural responses under anesthesia carry important information that are highly correlated with sensory perception. In the present study the usage of Urethane was chosen to minimize any effect from anesthetics (Capsius and Leppelsack, 1996; Curto et al., 2009; Hara and Harris, 2002). Furthermore, our findings suggesting distinct neural traces for Ref sequences over the AC in rats under anesthesia are comparable to the findings observed from human neuroimaging study during REM sleep (Andrillon et al., 2017). Thus, the present findings provide important initial findings on neural correlates during such passive, implicit learning in AC. Certain discrepancies between the present findings and previous human studies (e.g., no significant difference found from WN presentations) could be further studied by conducting experiments in awake animals.

In summary, the present study showed distinctive neural traits for re-occurring abstract auditory patterns that provide salient acoustic features (DRC) in the auditory cortex. While decreased induced activity amplitudes in the beta band observed throughout the auditory cortex suggests that both A1 and non-A1 areas are involved in encoding the information of re-occurring acoustic stimuli, such memory encoding in non-A1 areas could be processed in a shorter time frame. In this study we report, for the first time, a neural correlate of this type of memory formation in an easy to use, passive listening animal model, which should greatly facilitate further investigation into underlying neural mechanisms.

## Author Contributions

HK, RA, NA, MLS, and JS designed the study and ran pilots, HK and HA conducted the experiments, HK and RA analyzed the data, HK wrote the first draft of the manuscript, all authors revised the manuscript.

## References

Agus, T. R., Thorpe, S. J., and Pressnitzer, D. (2010). Rapid Formation of Robust Auditory Memories: Insights from Noise. Neuron, 66(4), 610–618. http://doi.org/10.1016/j.neuron.2010.04.014

Anderson, L. A., Christianson, G. B., and Linden, J. F. (2009). Stimulus-Specific Adaptation Occurs in the Auditory Thalamus. Journal of Neuroscience 29, 7359–7363. http://doi.org/10.1523/JNEUROSCI.0793-09.2009

Andrillon, T., Kouider, S., Agus, T., and Pressnitzer, D. (2015). Perceptual learning of acoustic noise generates memory-evoked potentials. Current Biology 25, 2823–2829. http://doi.org/10.1016/j.cub.2015.09.027

Andrillon, T., Pressnitzer, D., Léger, D., and Kouider, S. (2017). Formation and suppression of acoustic memories during human sleep. Nature Communications 8, 1–15. http://doi.org/10.1038/s41467-017-00071-z

Astikainen, P., Stefanics, G., Nokia, M., Lipponen, A., Cong, F., Penttonen, M., and Ruusuvirta, T. (2011). Memory-Based Mismatch Response to Frequency Changes in Rats. PLoS ONE 6, e24208–6. http://doi.org/10.1371/journal.pone.0024208

Auksztulewicz, R., and Friston, K. (2016). Repetition suppression and its contextual determinants in predictive coding. Cortex 80, 125–140. http://doi.org/10.1016/j.cortex.2015.11.024

Auksztulewicz, R., Friston, K. J., and Nobre, A. C. (2017). Task relevance modulates the behavioural and neural effects of sensory predictions. PLOS Biology 15, e2003143–27. http://doi.org/10.1371/journal.pbio.2003143

Ayala, Y. A., and Malmierca, M. S. (2013). Stimulus-specific adaptation and deviance detection in the inferior colliculus. Frontiers in Neural Circuits 6, 1–16. http://doi.org/10.3389/fncir.2012.00089/abstract

Bao, S., Chang, E. F., Woods, J., and Merzenich, M. M. (2004). Temporal plasticity in the primary auditory cortex induced by operant perceptual learning. Nature Neuroscience 7, 974–981. http://doi.org/10.1038/nn1293

Bar-Yosef, O., Rotman, Y., and Nelken, I. (2002). Responses of neurons in cat primary auditory cortex to bird chirps: Effects of temporal and spectral context. Journal of Neuroscience 22, 8619–8632.

Billig, A. J., Herrmann, B., Rhone, A. E., Gander, P. E., Nourski, K. V., Snoad, B. F., et al. (2019). A Sound-Sensitive Source of Alpha Oscillations in Human Non-Primary Auditory Cortex. Journal of Neuroscience 39, 8679–8689. http://doi.org/10.1523/JNEUROSCI.0696-19.2019

Capsius, B., and Leppelsack, H. (1996). Influence of urethane anesthesia on neural processing in the auditory cortex analogue of a songbird. Neuroscience Research 96, 59–70. http://doi.org/10.1016/0378-5955(96)00038-X

Curto, C., Sakata, S., Marguet, S., Itskov, V., and Harris, K. D. (2009). A Simple Model of Cortical Dynamics Explains Variability and State Dependence of Sensory Responses in Ure-thane-Anesthetized Auditory Cortex. Journal of Neuroscience 29, 10600–10612. http://doi.org/10.1523/JNEUROSCI.2053-09.2009

David, O., Kilner, J. M., and Friston, K. J. (2006). Mechanisms of evoked and induced responses in MEG/EEG. NeuroImage 31, 1580–1591. http://doi.org/10.1016/j.neuroimage.2006.02.034

de Cheveigné, A., and Arzounian, D. (2018). Robust detrending, rereferencing, outlier detection, and inpainting for multichannel data. NeuroImage 172, 903–912. http://doi.org/10.1016/j.neuroimage.2018.01.035

Gaese, B. H., and Ostwald, J. (2001). Anesthesia changes frequency tuning of neurons in the rat primary auditory cortex. Journal of Neurophysiology 86, 1062–1066. http://doi.org/10.1152/jn.2001.86.2.1062

Garrido, M. I., Kilner, J. M., Kiebel, S. J., Stephan, K. E., Baldeweg, T., and Friston, K. J. (2009). Repetition suppression and plasticity in the human brain. NeuroImage 48, 269–279. http://doi.org/10.1016/j.neuroimage.2009.06.034

Haenschel, C., Baldeweg, T., Croft, R. J., Whittington, M., and Gruzelier, J. (2000). Gamma and beta frequency oscillations in response to novel auditory stimuli: A comparison of human electroencephalogram (EEG) data with in vitro models. Proceedings of the National Academy of Sciences 97, 7645–7650. http://doi.org/10.1073/pnas.120162397

Hara, K., and Harris, R. A. (2002). The Anesthetic Mechanism of Urethane: The Effects on Neurotransmitter-Gated Ion Channels. Anesthesia and Analgesia 94, 313–318. http://doi.org/10.1213/00000539-200202000-00015

Hartmann, T., Schlee, W., and Weisz, N. (2012). It’s only in your head: expectancy of aversive auditory stimulation modulates stimulus-induced auditory cortical alpha desynchronization. NeuroImage 60, 170–178. http://doi.org/10.1016/j.neuroimage.2011.12.034

Huetz, C., Philibert, B., and Edeline, J. M. (2009). A Spike-Timing Code for Discriminating Conspecific Vocalizations in the Thalamocortical System of Anesthetized and Awake Guinea Pigs. Journal of Neuroscience 29, 334–350. http://doi.org/10.1523/JNEUROSCI.3269-08.2009

Klimesch, W., Russegger, H., Doppelmayr, M., and Pachinger, T. (1998). A method for the calculation of induced band power: implications for the significance of brain oscillations. Electroencephalography and Clinical Neurophysiology 108, 123–130. http://doi.org/10.1016/S0168-5597(97)00078-6

Lim, Y., Lagoy, R., Shinn-Cunningham, B. G., and Gardner, T. J. (2016). Transformation of temporal sequences in the zebra finch auditory system. eLife 5, e18205–18. http://doi.org/10.7554/eLife.18205

Lu, K., Liu, W., Zan, P., David, S. V., Fritz, J. B., and Shamma, S. A. (2018). Implicit Memory for Complex Sounds in Higher Auditory Cortex of the Ferret. Journal of Neuroscience 38, 9955–9966. http://doi.org/10.1523/JNEUROSCI.2118-18.2018

Luo, H., Tian, X., Song, K., Zhou, K., and Poeppel, D. (2013). Neural Response Phase Tracks How Listeners Learn New Acoustic Representations. Curbio 23, 968–974. http://doi.org/10.1016/j.cub.2013.04.031

Malmierca, M. S., Sanchez-Vives, M. V., Escera, C., and Bendixen, A. (2014). Neuronal adaptation, novelty detection and regularity encoding in audition. Frontiers in Systems Neuroscience 8, 1–9. http://doi.org/10.3389/fnsys.2014.00111

Nieto-Diego, J., and Malmierca, M. S. (2016). Topographic Distribution of Stimulus-Specific Adaptation across Auditory Cortical Fields in the Anesthetized Rat. PLOS Biology 14, e1002397–30. http://doi.org/10.1371/journal.pbio.1002397

Noda, T., and Takahashi, H. (2015). Anesthetic effects of isoflurane on the tonotopic map and neuronal population activity in the rat auditory cortex. European Journal of Neuroscience 42, 2298–2311. http://doi.org/10.1111/ejn.13007

Pagliardini, S., Greer, J. J., Funk, G. D., and Dickson, C. T. (2012). State-Dependent Modulation of Breathing in Urethane-Anesthetized Rats. Journal of Neuroscience 32, 11259–11270. http://doi.org/10.1523/JNEUROSCI.0948-12.2012

Parras, G. G., Nieto-Diego, J., Carbajal, G. V., Valdés-Baizabal, C., Escera, C., and Malmierca, M. S. (2017). Neurons along the auditory pathway exhibit a hierarchical organization of prediction error. Nature Communications 8, 1–17. http://doi.org/10.1038/s41467-017-02038-6

Pearce, M. T., Ruiz, M. H., Kapasi, S., Wiggins, G. A., and Bhattacharya, J. (2010). Unsupervised statistical learning underpins computational, behavioural, and neural manifestations of musical expectation. NeuroImage 50, 302–313. http://doi.org/10.1016/j.neuroimage.2009.12.019

Scholz, S., Schneider, S. L., and Rose, M. (2017). Differential effects of ongoing EEG beta and theta power on memory formation. PLoS ONE 12, e0171913–18. http://doi.org/10.1371/journal.pone.0171913

Skipper, J. I. (2014). Echoes of the spoken past: how auditory cortex hears context during speech perception. Philosophical Transactions of the Royal Society B: Biological Sciences 369, 20130297–19. http://doi.org/10.1098/rstb.2013.0297

Ulanovsky, N., Las, L., and Nelken, I. (2003). Processing of low-probability sounds by cortical neurons. Nature Neuroscience 6, 391–398. http://doi.org/10.1038/nn1032

Woods, V., Trumpis, M., Bent, B., Palopoli-Trojani, K., Chiang, C.-H., Wang, C., et al. (2018). Long-term recording reliability of liquid crystal polymer µECoG arrays. Journal of Neural Engineering 15, 066024–16. http://doi.org/10.1088/1741-2552/aae39d

Zurita, P., Villa, A. E. P., de Ribaupierre, Y., de Ribaupierre, F., and Rouiller, E. M. (1994). Changes of single unit activity in the cat’s auditory thalamus and cortex associated to different anesthetic conditions. Neuroscience Research 19, 303–316. http://doi.org/10.1016/0168-0102(94)90043-4

